# *Salmonella*-vectored vaccine delivering three *Clostridium perfringens* antigens protects poultry against necrotic enteritis

**DOI:** 10.1101/318469

**Authors:** Shyra Wilde, Yanlong Jiang, Amanda M. Tafoya, Jamie Horsman, Miranda Yousif, Luis Armando Vazquez, Kenneth L. Roland

## Abstract

Necrotic enteritis is an economically important poultry disease caused by the bacterium *Clostridium perfringens*. There are currently no necrotic enteritis vaccines available for use in broiler birds, the most important target population. *Salmonella*-vectored vaccines represent a convenient and effective option for controlling this disease. We used a single attenuated *Salmonella* vaccine strain, engineered to lyse within the host, to deliver up to three *C. perfringens* antigens. Two of the antigens were toxoids, based on *C. perfringens* α-toxin and NetB toxin. The third antigen was fructose-1,6-bisphosphate aldolase (Fba), an metabolic enzyme with an unknown role in virulence. Oral immunization with a single *Salmonella* vaccine strain producing either Fba, α-toxoid and NetB toxoid, or all three antigens, was immunogenic, inducing serum, cellular and mucosal responses against *Salmonella* and the vectored *C. perfringens* antigens. All three vaccine strains were protective against virulent *C. perfringens* challenge. The strains delivering Fba only or all three antigens provided the best protection. We also demonstrate that both toxins and Fba are present on the *C. perfringens* cell surface. The presence of Fba on the cell surface suggests that Fba may function as an adhesin.

## Introduction

Keeping our food supply safe is one of many challenges facing the agriculture industry. When rearing poultry, maintenance of a healthy flock requires a multi-faceted strategy that includes good husbandry, strong biosecurity practices, proper feed formulation, vaccination, and veterinary care. One element of poultry rearing has been the inclusion of sub-clinical amounts of antibiotics in the feed to promote growth. Recent concerns regarding the impact of this practice on increasing antibiotic resistance in human pathogens has led to stricter regulations governing antibiotic use in food animals and voluntary elimination of antibiotics by poultry producers and food providers. While limiting the use of antibiotics on the farm may have long-range health benefits for the human population, it leads to additional challenges for the poultry industry. Based on results in other countries, it is well known that the incidence of necrotic enteritis (NE) caused by *Clostridium perfringens* type A strains increases when antibiotics are removed from the feed [1].

NE is an enteric disease causing chronic mucosal damage to the intestines with a range of symptoms including general poor health, reduced appetite, reduced weight gain, poor digestion and cholangiohepatitis. More severe symptoms, such as sudden death, can also occur in afflicted flocks. There are often no overt symptoms of pathology. Subacute infections are the most common, resulting in economic losses, due to the reduced weight of the birds, and carcass condemnation, due to liver lesions, after slaughter [2]. There are many predisposing factors, which include feed composition, stress, coccidiosis, and immunosuppression due to infection with certain viruses [3]. This disease is estimated to cause annual global losses of up to $6 million dollars to poultry producers.

Vaccination is one practical alternative to antibiotics. In a previous report, we used a novel *Salmonella* Typhimurium vaccine vector to deliver two relevant clostridial toxoid antigens, PlcC, a nontoxic carboxyterminal fragment of α-toxin, and a GST-NetB fusion protein [4]. NetB is a pore-forming toxin that plays a central role in NE [5]. Although the role of α-toxin in pathogenesis is not clear, anti-α-toxin antibodies are protective [6], possibly due to their ability to inhibit *C. perfringens* growth [7]. The *S*. Typhimurium vaccine strain we used was engineered to display a near wild-type phenotype at the time of immunization. After several rounds of replication in host tissues, the strain lyses, releasing the *C. perfringens* antigens. This type of attenuated strain is called a lysis strain [8]. Immunization with the lysis strain delivering PlcC and GST-NetB was previously shown to elicit protective immunity against *C. perfringens* challenge [4].

Fructose-1,6-bisphosphate aldolase (Fba) was previously identified as an immunogenic protein in the supernatant of a virulent NE strain [9]. Chickens injected with recombinant Fba were partially protected against NE after challenge with *C. perfringens* [10]. In this work, we provide evidence that Fba is present on the surface of *C. perfringens* and evaluate the efficacy of Fba delivered by a *Salmonella* lysis strain with or without co-delivery of PlcC and NetB. We demonstrate that all three antigens can be effectively delivered by a single *Salmonella* lysis strain, eliciting strong mucosal and cellular responses. Inclusion of Fba enhanced protection against NE.

## Results

### Characterization of *Salmonella* vaccine strains

All *S*. Typhimurium vaccine strains required arabinose for growth (data not shown) and had similar growth characteristics in LB broth (0.1% arabinose, 0.1% mannose) (**Fig. S1**). Production of PlcC, GST-NetB and Fba by the various *Salmonella* vaccine strains were assessed by western blot. The strain carrying pYA5130 (*plcC fba* GST-*netB*) produced Fba at levels similar to the strain producing Fba alone (**Fig. 1**) and produced GST-NetB at levels similar to the strain carrying pYA5112 (*plcC* GST-*netB*). The strain carrying pYA5112 (*plcC* GST-*netB*)produced about 6-fold more PlcC than the strain carrying pYA5130 (*plcC fba* GST-*netB*) (**Fig. 1**). This is not surprising, as the *plcC* leader sequence in pYA5112 was modified to increase PlcC synthesis [4], while the *plcC* gene in pYA5130 does not carry that modification.

**Figure 1.**
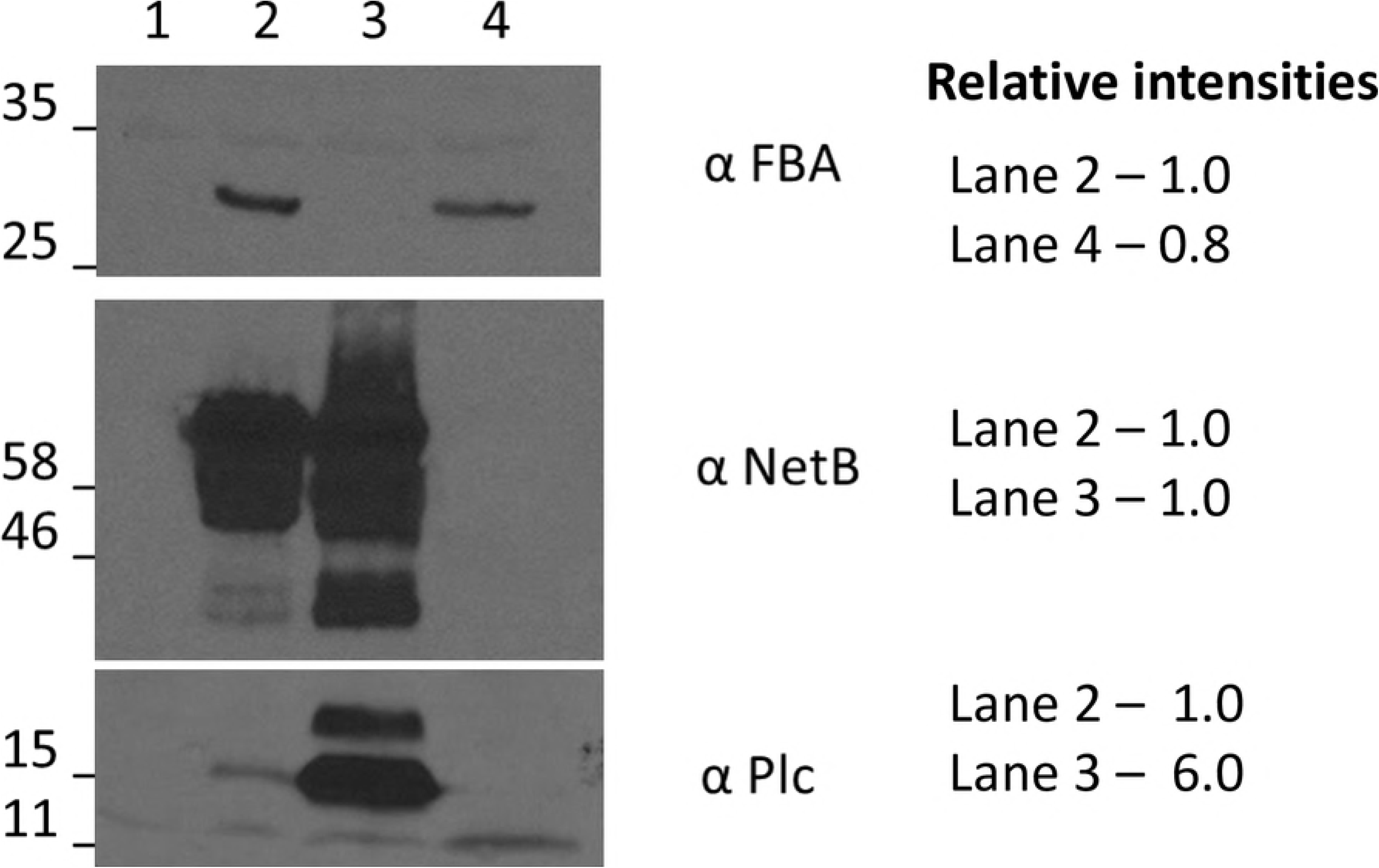
Antigen production by vaccine strains as determined by western blot. Vaccine strains were grown in LB as described in Materials and Methods. Antigen production was induced by the addition of IPTG four hours prior to harvest. Membranes were probed with the indicated anti-sera. Predicted mass of antigens are: PlcC, 18kDa; GST-NetB, 59 kDa; Fba, 30 kDa. Lane 1, *Salmonella* vector control; Lane 2, *Salmonella* strain carrying pYA5130 (*plcC, fba, netB*); Lane 3, *Salmonella* strain carrying pYA5112 (*plcC*, *netB*); Lane 4, *Salmonella* strain carrying pKR023 (*fba*).

### Serum antibody responses

In experiment 1, the triple antigen vaccine elicited a significant increase in anti-Fba serum IgY titers when compared to the double-antigen vaccine or vector control (*P* < 0.0001) (**Fig. 2A**). Modest increases in anti-PlcC and anti-NetB responses were observed, although only the anti-NetB responses elicited by the triple antigen vaccine achieved significance (*P* < 0.02). In experiments 2 and 3, birds received 10-fold higher vaccine doses. Results from experiments 2 and 3 were similar and the data were combined for analysis. In this case, the anti-Fba serum IgY titers were 14 – 21-fold greater than controls, which was significant (**Fig. 2B**; *P* < 0.02). We observed similar, significant increases in anti-NetB titers in birds immunized with *Salmonella* vaccines producing NetB (*P* < 0.05). Despite the difference in PlcC production by strains carrying pYA5112 and pYA5130 (Fig. 1), the serum responses elicited by the two strains were comparable in both experiments (Fig. 2). However, though we observed increased anti-PlcC titers in birds immunized with strains producing PlcC, the increases were not statistically significant. This result is consistent with results from our previous study in which we observed poor seroconversion to PlcC when delivered by the same double antigen strain [4]. The anti-*Salmonella* LPS titers in immunized birds were 8 – 32-fold higher than controls (*P* < 0.0001) (**Fig. 2B**).

**Figure 2.**
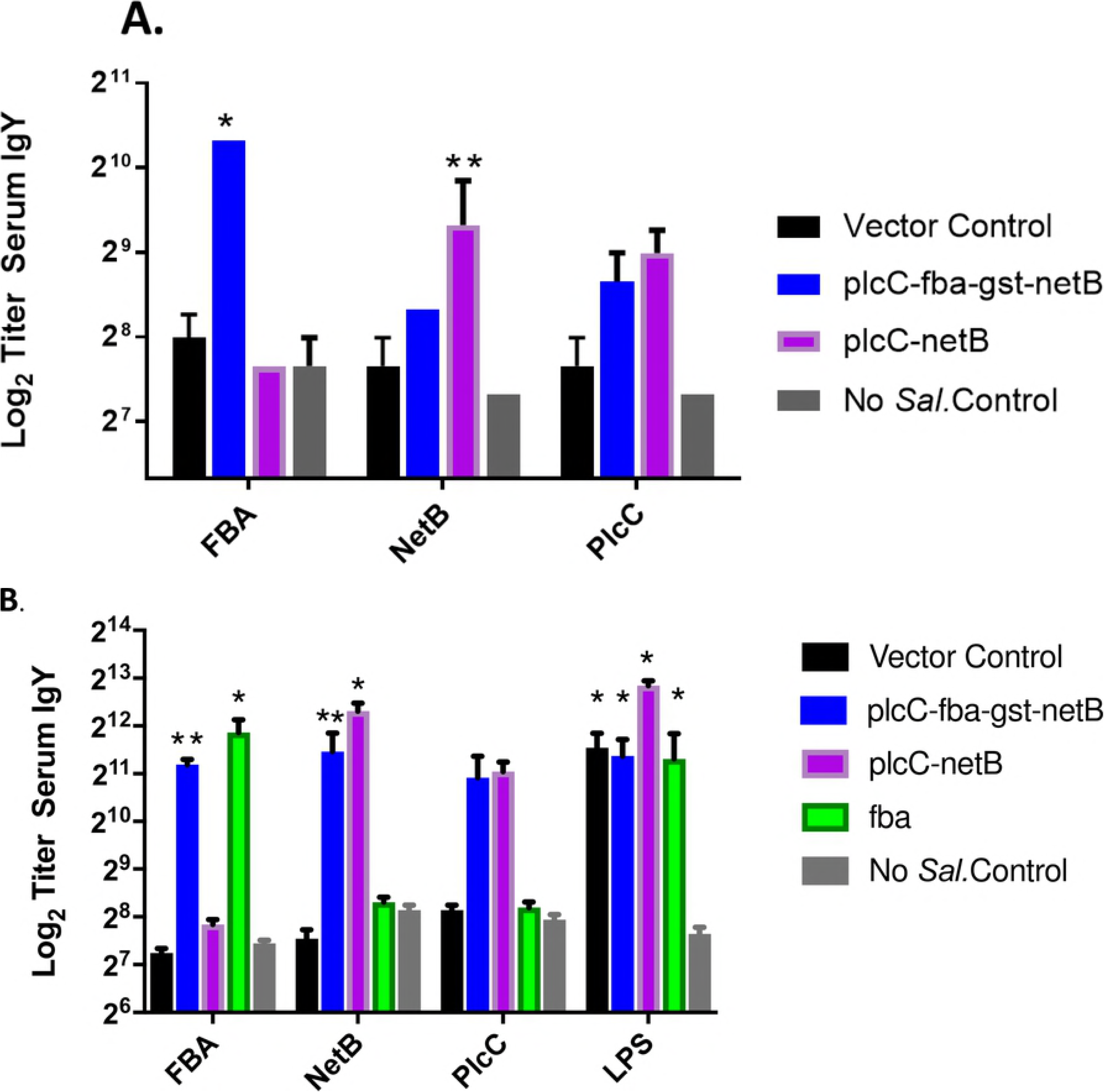
Serum antibodies against *C. perfringens* and *Salmonella* antigens in vaccinated and non-vaccinated birds as determined by ELISA. (A) Experiment 1; (B) Experiments 2/3. Differences in responses compared to non-vaccinates and vector-only controls are indicated *, *P* < 0.0001; **, *P* < 0.02. LPS responses were compared to non-vaccinated controls.

### Mucosal IgA, IgM and IgY responses

We examined intestinal mucosal responses in experiments 2 and 3 and have combined the data. The vaccine elicited strong, significant mucosal responses to all antigens, including PlcC, although statistically significant responses against every antigen were not observed for all isotypes (**Fig. 3**). The mucosal IgA responses against PlcC were not significant for either strain delivering PlcC, while mucosal IgM responses elicited by both strains were significant. The anti-PlcC mucosal IgY response was only significant for the strain delivering two antigens.

**Figure 3.**
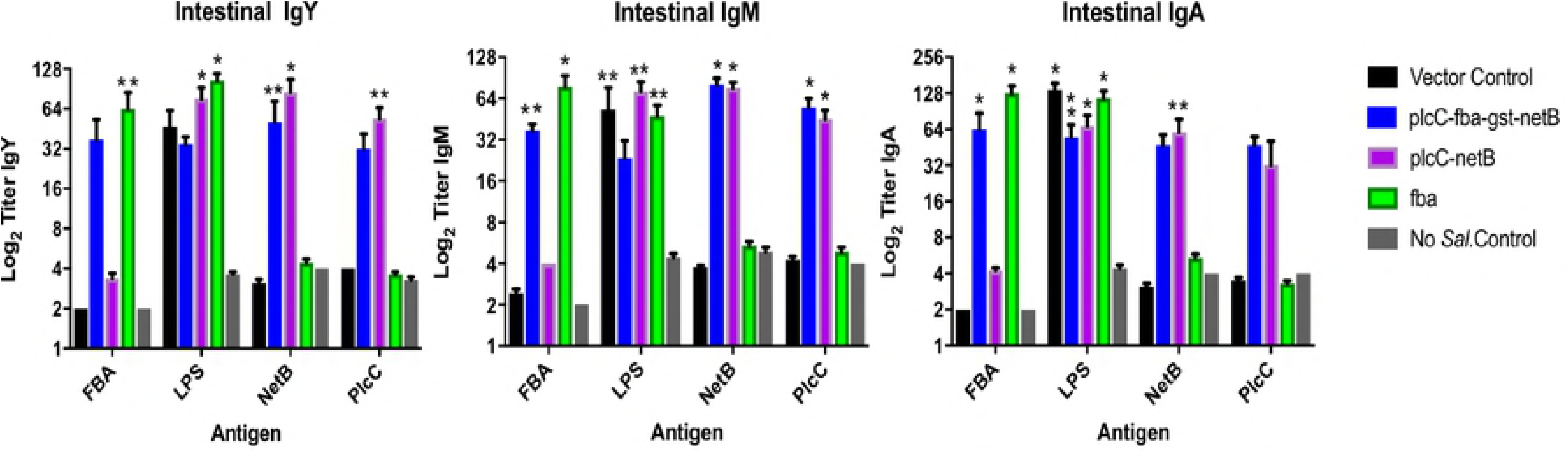
Mucosal antibody responses to the indicated *C. perfringens* proteins and *Salmonella* LPS as determined by ELISA. (A) Experiment 1; (B) Experiments 2/3. Differences in responses compared to non-vaccinates and vector-only controls are indicated by *, *P* < 0.0001; **, *P* < 0.02. LPS responses were compared to non-vaccinated controls.

Birds immunized with the strain delivering only Fba achieved significant anti-Fba mucosal responses across all isotypes (*P* < 0.02). The anti-Fba IgM and IgA responses were significant in birds immunized with the triple antigen strain. The anti-Fba IgY responses, while elevated, were not significant (*P* = 0.20). All immunized birds produced elevated anti-NetB responses, although the anti-NetB IgA responses in birds that received the triple antigen strain did not achieve significance (*P* = 0.08)

### Cellular responses

Immunization with the vaccine producing all three antigens elicited strong anti-Fba and anti-PlcC cellular responses in blood lymphocytes and splenocytes (**Fig. 4**). Significant anti-NetB responses were observed in the splenocytes of birds receiving the triple antigen strain (*P* < 0.05), but not in lymphocytes. Birds immunized with the strain producing two antigens, PlcC and NetB, produced significant anti-NetB responses in splenocytes (*P* < 0.005) and significant anti-PlcC responses in lymphocytes (*P* < 0.05). Birds immunized with the strain producing Fba alone exhibited significant cellular responses in splenocytes (*P* < 0.0001), while the response in lymphocytes was not significant.

**Figure 4.**
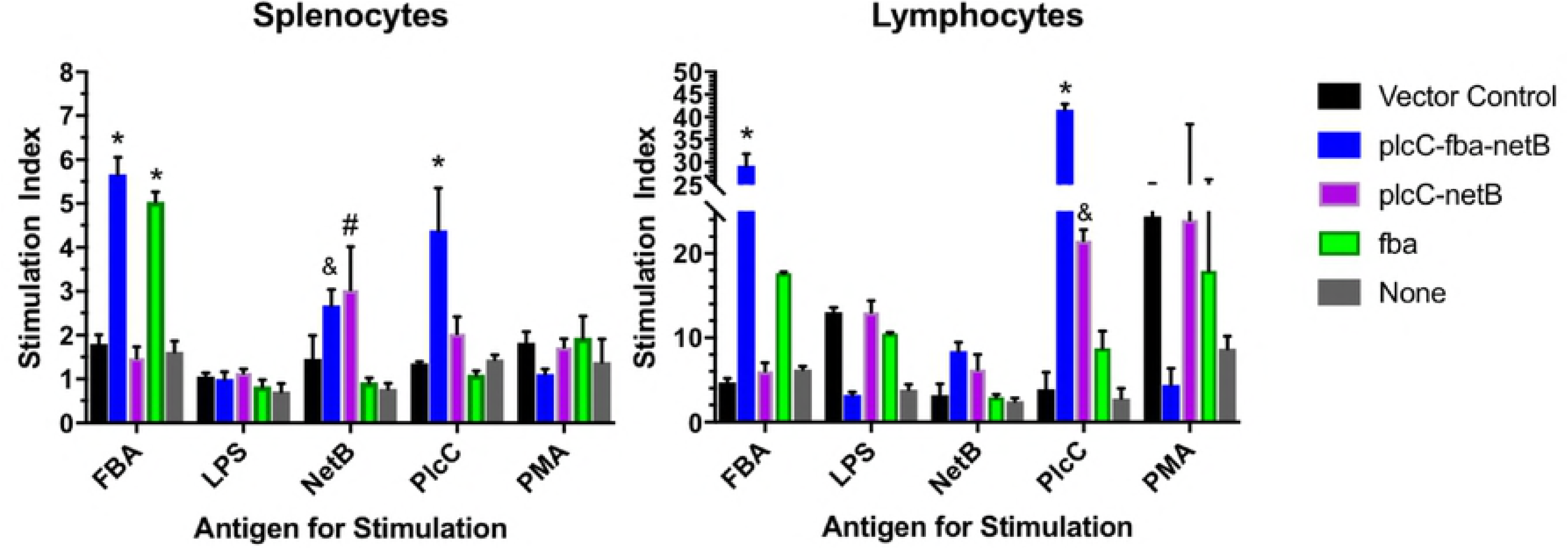
Cellular responses by splenocytes and lymphocytes from vaccinated and non-vaccinated chickens. Significant differences compared to controls are indicated. ^*^, *P* < 0.0001; ^#^, *P* < 0.005, ^&^, *P* < 0.05.

### Protection studies

Protection against challenge was assessed by scoring intestinal lesions using a 6 point scoring system [11]. In experiment 1, birds immunized with either the double antigen strain or the triple antigen strain had significantly lower lesion scores than birds in the control groups (**Table 1**). There was no statistical difference between the average lesion scores for the triple antigen and double antigen strain groups (*P* = 0.25) in this experiment.

**Table 1.**
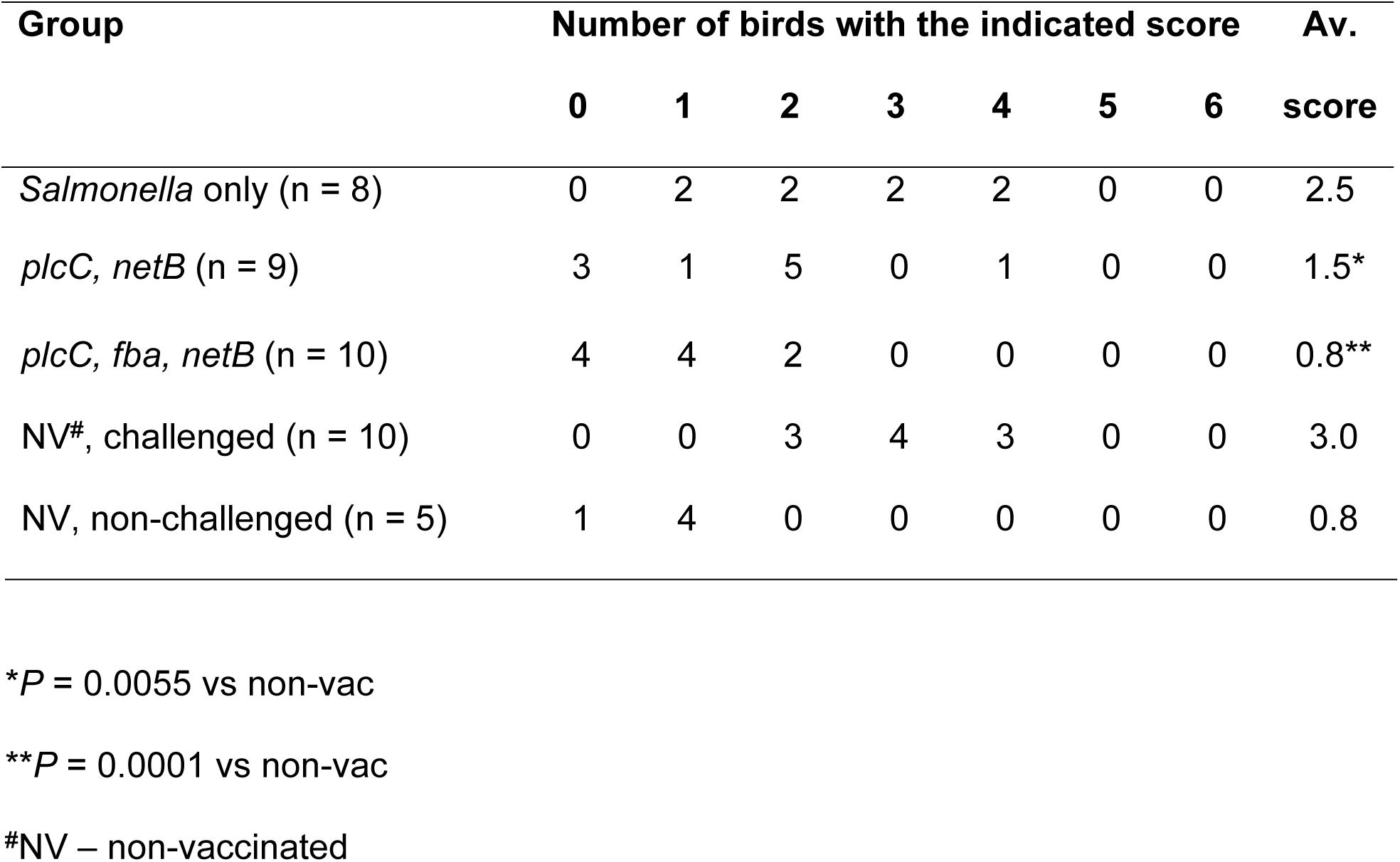
Intestinal lesion scores from Experiment 1

In experiment 3, we included a strain producing only Fba in addition to the strains evaluated in experiment 1. Chickens immunized with any strain carrying *C. perfringens* antigens had significantly lower lesion scores than non-vaccinated or *Salmonella* only controls (**Table 2**), indicating protection against *C. perfringens* challenge. As in the previous experiment, there were no statistical differences between the double antigen and triple antigen groups (*P* = 0.19). However, there was a significant difference in lesion scores between the Fba-only group and the double antigen group (*P* = 0.01), indicating that Fba is contributing to protection against *C. perfringens* challenge.

**Table 2.** Intestinal lesion scores from Experiment 3

| Group | Number of birds with the indicated score |  |  |  |  |  |  | Av. |
| --- | --- | --- | --- | --- | --- | --- | --- | --- |
|  | 0 | 1 | 2 | 3 | 4 | 5 | 6 | score |
| *Salmonella* only (n = 15) | 0 | 1 | 3 | 7 | 2 | 2 | 0 | 3.1 |
| *plcC, netB* (n = 14) | 2 | 4 | 5 | 2 | 0 | 0 | 1 | 1.9* |
| *plcC, fba, netB* (n = 17) | 5 | 6 | 4 | 2 | 0 | 0 | 0 | 0.9** |
| *fba* (n = 16) | 8 | 5 | 3 | 0 | 0 | 0 | 0 | 0.7**& |
| NV, challenged (n = 9) | 0 | 0 | 1 | 2 | 4 | 0 | 2 | 4.0 |
| NV, non-challenged (n = 6) | 3 | 3 | 0 | 0 | 0 | 0 | 0 | 0.5 |

\* $P = 0.0012$ vs non-vac, $P = 0.0056$ vs *Salmonella* only group
\*\* $P < 0.0001$ vs non-vac or *Salmonella* only groups
& $P = 0.01$ vs *plcC, netB* group

There were no significant differences between any vaccinated group and the non-challenged controls.

### Plc, NetB and Fba are displayed on the surface of *C. perfringens*

To evaluate the potential of rabbit antisera raised against rPlcC, rNetB and rFba to bind directly to *C. perfringens*, we used an immunofluorescence assay. Previous work showed that anti-PlcC antibodies bind to the surface of *C. perfringens* [7]. We confirmed this observation. Rabbit antibodies raised against rPlcC bound to challenge strain CP4 and another NE strain, JGS4143 (**Fig. 5**). Binding was dependent on the presence of α-toxin, as no fluorescence was observed when we probed JGS5388, a ∆*plc* derivative of JGS4143. In addition, when purified rPlcC was mixed with the antisera prior to incubation with CP4, immunofluorescence was drastically reduced, indicating that fluorescence was mediated by anti-α-toxin-specific antibodies present in the serum. When we probed wild-type *C. perfringens* strains CP4 and JGS4143 with anti-NetB antisera, we observed immunofluorescence. We also observed immunofluorescence using ∆*plc* strain JGS5388. However, no immunofluorescence was observed using the naturally occurring NetB-strain JGS4043 or when we mixed purified rNetB with the anti-NetB antisera prior to incubation with CP4 (**Fig. 5,** right column).

**Figure 5.**
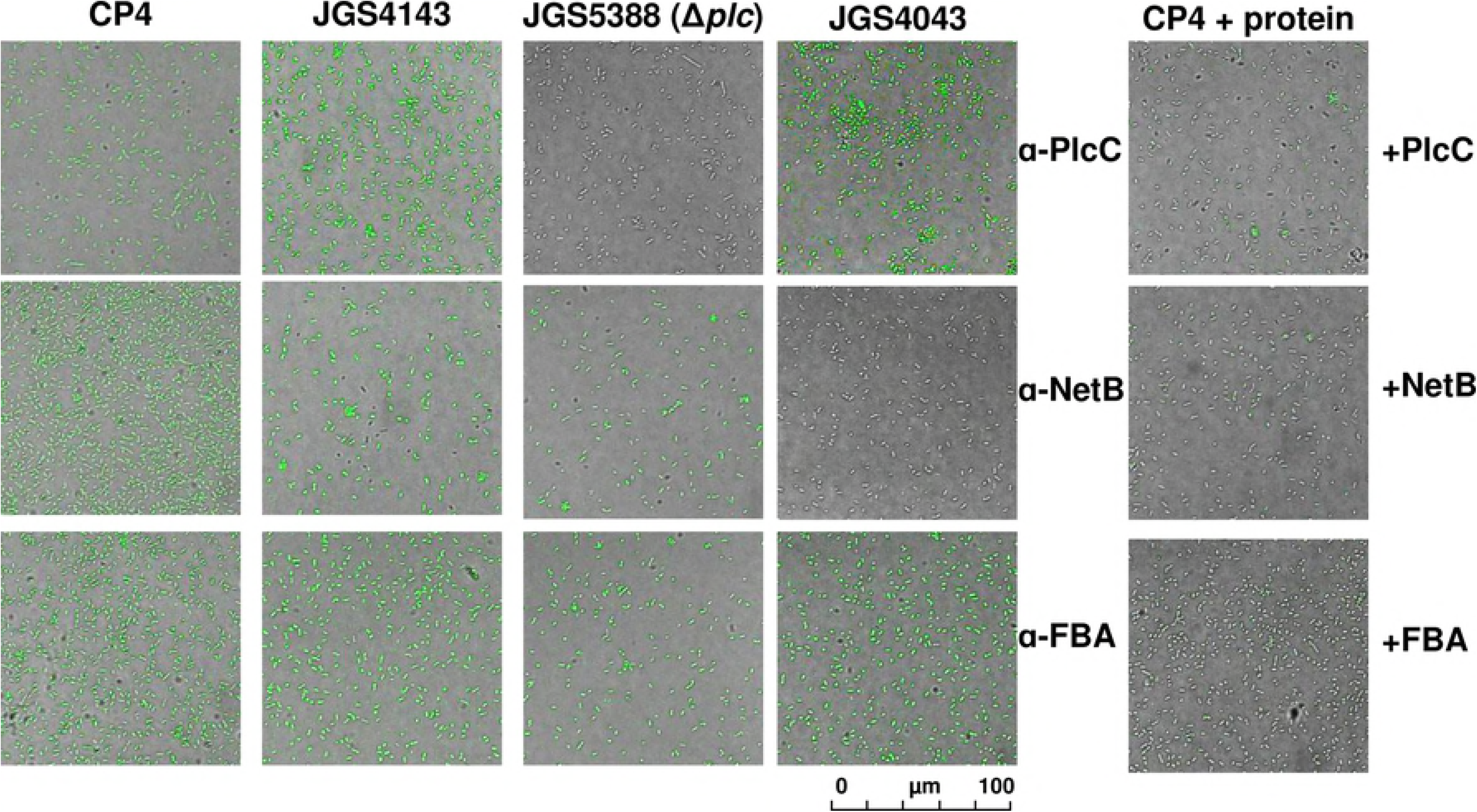
Indirect immunofluorescence detection of proteins on the surface of *C. perfringens*. *C. perfringens* strains were incubated with polyclonal sera from rabbits hyperimmunized with the indicated *C. perfringens* proteins. In the far right panel, sera were mixed with 1 μg of the indicated proteins for 30 min prior to incubation with the *C. perfringens* strains.

Interestingly, Fba appears to be present on the cell surface as well, since we observed immunofluorescence of all strains when probed with anti-Fba antisera. Fluorescence was lost when the sera was pre-incubated with purified rFba (**Fig. 5**, right column). Incubation of *C. perfringens* with pre-immune sera or the secondary antibody alone did not result in immunofluorescence (**Fig. S2**). As an additional control, we incubated purified rFba with the anti-NetB antisera and rNetB with the anti-Fba antisera prior to probing *C. perfringens* CP4. These incubations did not diminish immunofluorescence (**Fig. S2**), indicating that protein-specific titration of each antibody was required to prevent its binding to *C. perfringens*. This result supports our interpretation that NetB and Fba are displayed on the surface of *C. perfringens*.

### NetB and Fba facilitate adherence to eukaryotic cells

The presence of Fba on the *C. perfringens* cell surface raises the possibility that it plays a role in attachment to the host epithelium. To investigate the possible role of Fba in adherence, we examined the effect of pre-incubating Caco-2 cells with Fba prior to performing an attachment assay. As controls, we included wells in which either NetB or the *Streptococcus pneumoniae* protein PspA was substituted for Fba. Prior addition of Fba to the wells resulted in a significant reduction in adherence (*P* < 0.037) (**Fig. 6**). We also found that NetB inhibited adherence as well (*P* < 0.002). The irrelevant *S. pneumoniae* protein PspA had no effect. These results suggest a possible role for Fba and NetB in adherence to intestinal epithelial cells.

**Figure 6.**
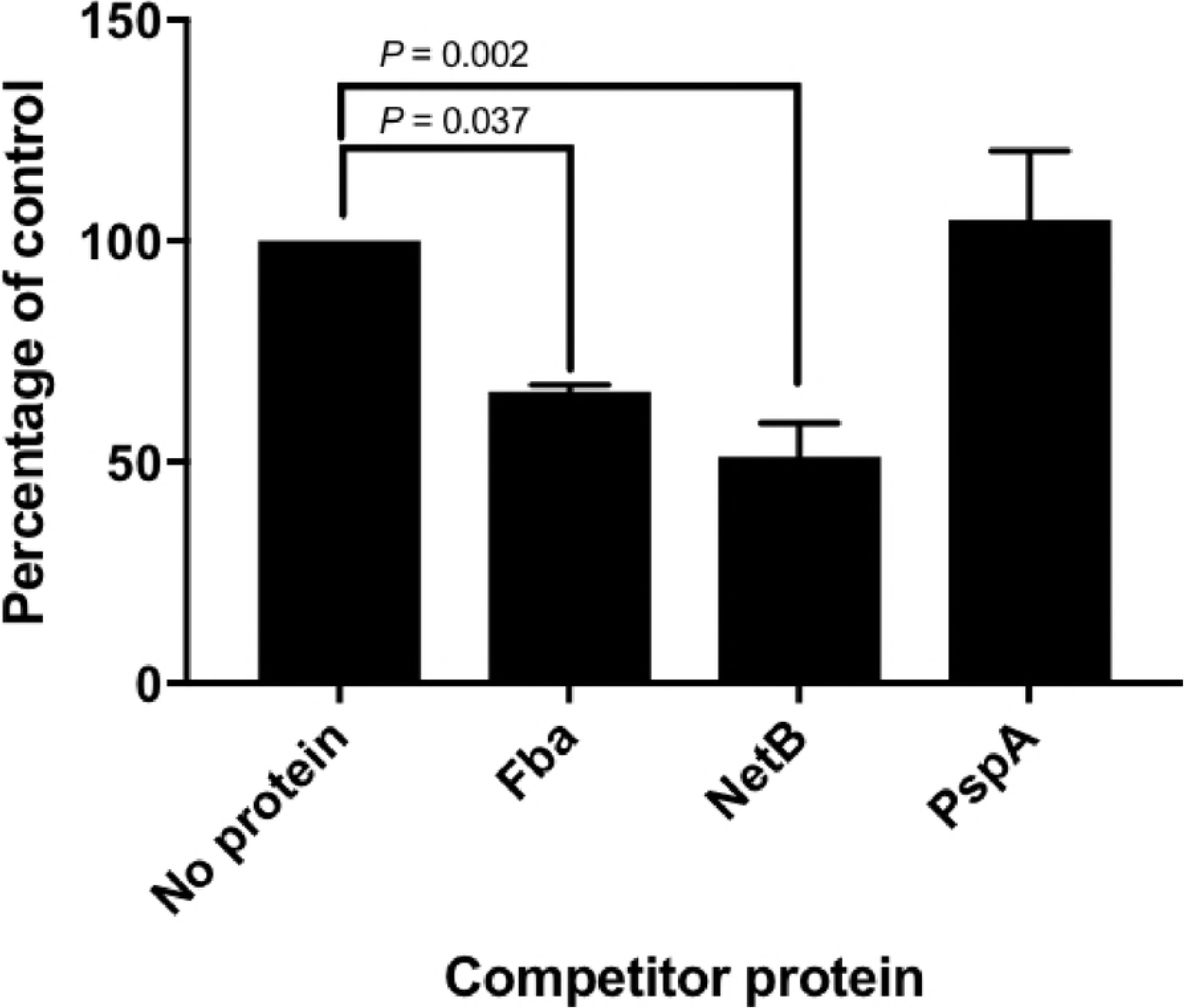
Adherence of *C. perfringens* CP4 to Caco-2 cells in the presence or absence of purified rFba, rNetB or rPspA. Attachment results significantly different from the no protein controls are indicated.

## Discussion

Fba is an enzyme of the Embden-Meyerhof-Parnas glycolytic pathway that converts D-fructose-1,6-bisphosphate into D-glyceraldehyde-3-phosphate and dihydroxyacetone phosphate [12]. In *C. perfringens*, it also plays a role in the catabolism of *myo*-inositol [13] and is negatively regulated by the VirR/VirS two component system that also regulates production of a number of virulence factors including α-toxin [14]. An enzyme important in central metabolism may seem to be an unusual choice for inclusion in a vaccine. However, in several organisms, Fba is recognized as a “moonlighting protein”, one that can perform two or more autonomous functions [12]. Fba has been shown to serve as an adhesin in a number of dissimilar bacterial pathogens, including *S. pneumoniae* [15], *Mycobacterium tuberculosis* [16] and *Neisseria meningitidis* [17]. Our immunofluorescence data shows that a protein cross-reactive with anti-Fba antisera is present on the surface of *C. perfringens* (**Fig. 5**), a result consistent with a possible role for Fba as an adhesin in this organism. Our tissue culture results support this as well, as adherence to Caco-2 cells is inhibited in the presence of rFba (**Fig. 6**). However, the importance of this finding will need to be confirmed using a chicken epithelial cell line. Similarly, NetB appears to aid in adherence to epithelial cells. This is not surprising, since NetB is known to be toxic to both chicken and human cells. NetB may bind to host cells via interactions with cholesterol [18].

As a vaccine, immunization with the appropriate homologous Fba is partially protective against infection with *S. pneumoniae* [19], *S. pyogenes* [20] and the fish pathogen, *Edwardsiella tarda* [21]. In *C. perfringens*, Fba was previously identified as one of several secreted proteins recognized by sera taken from chickens with acquired immunity to necrotic enteritis [9]. In an initial assessment of its vaccine potential, chickens given a single intramuscular injection with recombinant Fba were well protected against a mild *C. perfringens* challenge [10]. Protection against a more severe challenge required multiple injections. Fba delivered by a non-lysis attenuated *S*. Typhimurium vaccine was also found to elicit partially protective immunity [22], although protection was not as effective as when it was delivered by intramuscular injection. In the current study, Fba delivered by an attenuated lysis *S*. Typhimurium strain elicited the strongest protection, significantly better than the vaccine delivering the two toxoid antigens (**Table 2**). It is possible that, despite the careful strain design to prevent loss of immunogenicity due to antigen load, there is some stress on the vaccine strain delivering the two toxoid antigens, leading to lower immune responses against each antigen. However, in the triple antigen strain, this deficiency appears to be offset by the combined protective efficacy provided by each antigen. Although immunization with the strain delivering Fba alone yielded the lowest lesion scores, it is likely that the triple antigen strain will provide the broadest protection. This question will be examined in future experiments.

It is of interest that, in this study, we achieved anti-Fba serum IgY titers similar to the serum titers observed when Fba was delivered by a non-lysis strain [22]. Conversely, delivery by lysis strain elicited much greater mucosal responses, indicating that mucosal immunity is more important than humoral immunity for protection against challenge.

Previous work demonstrated that anti-PlcC antibodies bound to the surface of *C. perfringens*, indicating the presence of α-toxin [7]. We extended those observations to confirm that antibody binding did not occur in a ∆*plc* mutant and that binding is reduced by incubation of antisera with purified PlcC (**Fig. 5**). Our data also indicate that NetB toxin is present on the surface of *C. perfringens*. The presence of all three antigens on the cell surface suggests that the anti-*C. perfringens* mucosal antibodies elicited by our vaccine may bind directly to *C. perfringens*, facilitating opsonization and/or inhibiting toxin secretion. If, as our data suggest, Fba serves as an adhesin for *C. perfringens*, anti-Fba antibodies bound to *C. perfringens* cells may serve to prevent close contact of the bacterium with the host epithelium, an important step in pathogenesis [23, 24]. In addition, anti-PlcC antibodies were previously shown to inhibit *C. perfringens* growth directly, suggesting another mechanism for the action of these antibodies.

This work highlights the potential for a *Salmonella*-vectored vaccine to control NE. Our findings show that the *Salmonella* lysis vector strain χ11802 was capable of delivering up to three antigens simultaneously, generating humoral, cellular and mucosal responses against all antigens. In particular, this strain is able to generate strong mucosal responses as shown here and in a previous study [4]. Our results also support the idea that intestinal mucosal responses are an effective deterrent against lesion formation caused by *C. perfringens*.

## Materials and Methods

### Animal care

All animal experiments were conducted in compliance with the Arizona State University Institutional Animal Care and Use Committee and the Animal Welfare Act under protocol 16-1480R.

### Bacterial strains, plasmids and growth conditions

Strains and plasmids used in this study are listed in **Table 3**. All vaccine strains were routinely cultured at 37°C with aeration. *Salmonella* were grown in Luria Broth (LB) (Bacto tryptone, 10 g/liter; Bacto yeast extract, 5 g/liter; NaCl, 10 g/liter) with aeration. When required, media were supplemented with ampicillin (Amp; 100 μg/mL), 2,6-diaminopimelic acid (DAP; 50 μg/mL), L-arabinose (Ara; 0.1% v/v) or mannose (Man; 0.1% v/v). Media were solidified with 1.5% (wt/vol) agar as needed. Vaccine strains were supplemented with 0.1% Ara in plates. *C. perfringens* was cultured anaerobically on blood agar plates, in cooked meat medium (CMM; Difco) or in fluid thioglycollate medium (FTG; Difco) for challenge.

**Table 3.**
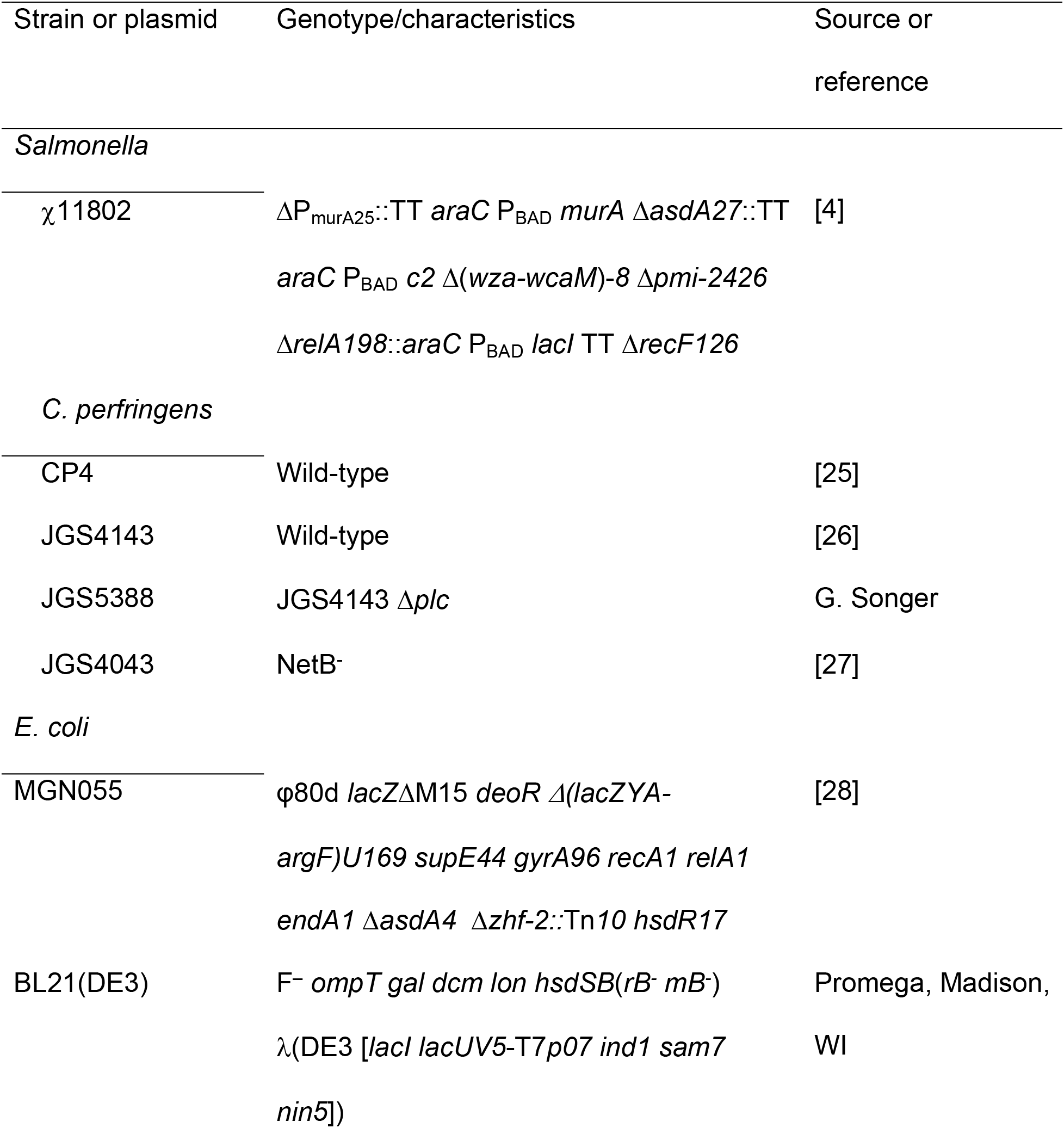
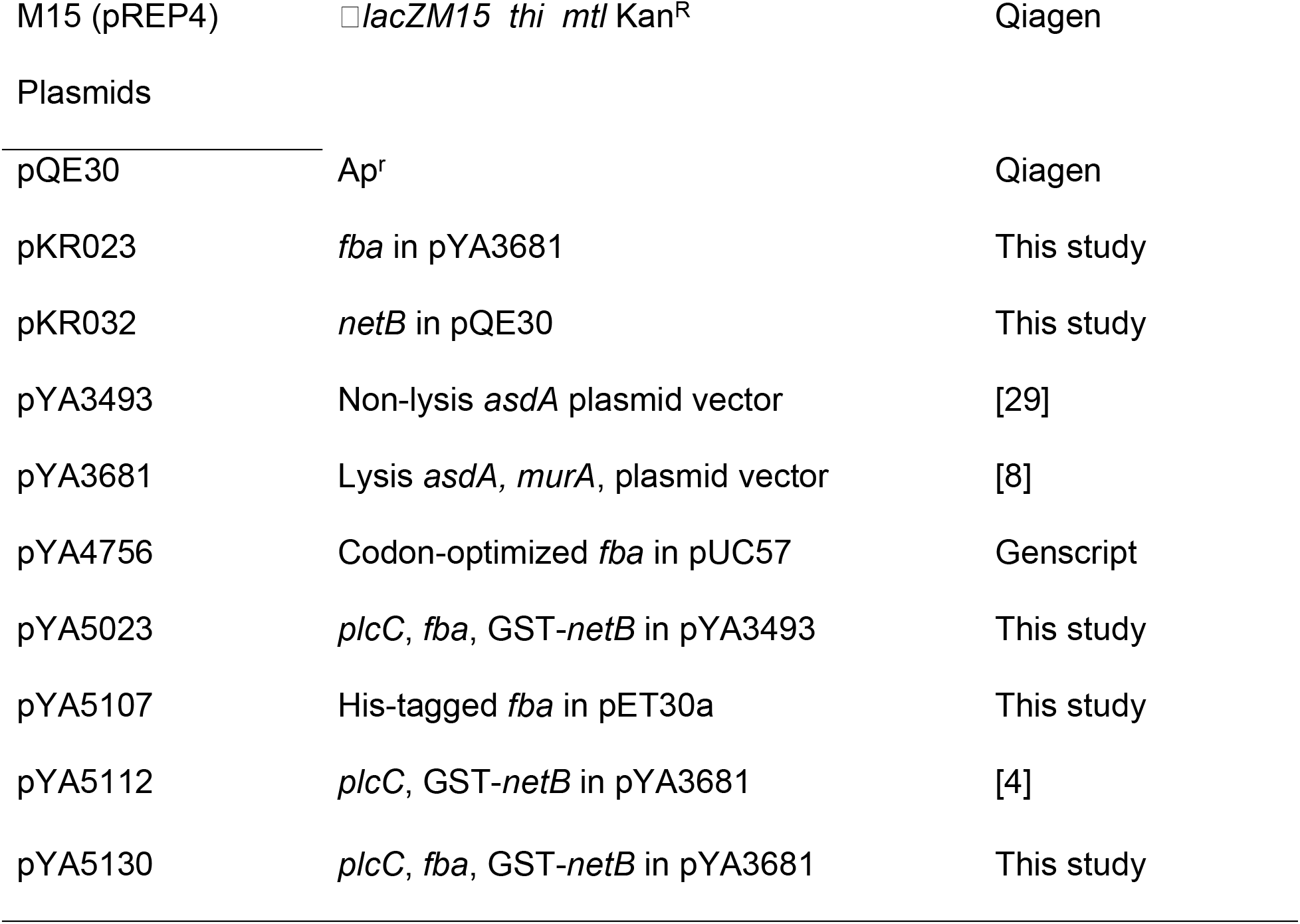
Strains and plasmids used in this study.

### Plasmid constructions

The *fba* gene, codon-optimized for expression in *Salmonella*, was synthesized by Genscript (Piscataway, NJ, USA). To construct plasmid pKR023, the *fba* gene was amplified from plasmid pYA4756 using primers Fba-KpnI-F: (TTGGGGTACCTCATGGCACTGGTTAACGCAAAAG) and Fba-SacII: ATTACCGCGGCTATTAAGCTCTGTTTACTGA. Primers 3681-KpnI (TGGGGTACCAGATGGCACTGGTTAACGCAAAAG) and 3681-SacII (ATTACCGCGGCTATTAAGCTCTGTTTACTGA) were used to introduce KpnI and SacII sites into pYA3681. The *fba* gene was cloned into pYA3681, electroporated into *Escherichia coli* strain MGN055 and then subsequently moved into *S*. Typhimurium strain χ11802. To construct plasmid pYA5130, primers 5023-KpnI-F (CCATGGGGTACCAGATGAGTATTCAACATTTCCGT) and 5023-SacII (ATTACCGCGGTTACAGATAATATTCGATTTTAATT) were used to amplify the *plcC-fba-Gst-netB* gene cassette from pYA5023. The purified PCR product was digested with KpnI and SacII. Primers 3681-KpnI and 3681-SacII were used to amplify the plasmid sequences from pYA3681. The purified PCR product was digested with KpnI and SacII. The two fragments were then ligated and electroporated into MGN055. A plasmid of the expected size and DNA sequence was designated pYA5130.

### Immunofluorescence assay

To determine if antibodies against NetB, PlcC, and Fba bound to the bacterial cell surface, we performed an indirect immunofluorescence test. Several fresh colonies of *C. perfringens* from a Trypticase Soy Agar with 5% Sheep Blood plate (BD) were used to inoculate Brain Heart Infusion (BHI) Broth (BD). Cultures were grown for 24 h and harvested by centrifugation at 4000 X *g* for 15 min. Pelleted cells were resuspended with 10% formalin in phosphate buffered saline (PBS) and fixed by overnight incubation at 4°C with slow rotation. Fixed cells were pelleted, resuspended, and washed twice in PBS. 100μL of fixed cells were pelleted and blocked with 2% BSA for 1 h at room temperature with slow agitation. Cells were pelleted and resuspended in 200 μL of control sera, anti-sera, anti-sera + antigen or control protein, or PBS (no sera control) diluted 1:50 in PBS and incubated overnight at 4°C with slow rotation. Cells were washed with PBS-0.1% Tween 20 and goat anti-chicken IgG antibody conjugated with fluorescein isothiocyanate (SouthernBiotech, Birmingham, AL) diluted 1:500 in PBS was added. The resulting cell suspension was incubated for 3 h at room temperature with gentle agitation. Cells were washed 3 times with PBS-0.1% Tween 20, resuspended in PBS and mounted for observation under a Leica TCS SP5 confocal microscope.

### Synthesis of recombinant antigens in χ11802

To evaluate synthesis of PlcC, Fba and GST-NetB, overnight static cultures of *S.* Typhimurium χ11802 carrying plasmids pYA5112 (PlcC, GST-NetB), pYA5130 (PlcC, Fba, GST-NetB), pKR029 (Fba) or pYA3681 (empty vector) were inoculated into LB supplemented with 0.1% arabinose and 0.2% mannose and grown to an OD_600_ of 0.6. Then, 1 mM isopropyl-β-D-thiogalactopyranoside (IPTG) was added to each culture and induced for an additional 4 h. The final cultures were adjusted to the same density using OD_600_ values. Equal volumes of the adjusted samples were centrifuged at 16,000 × *g* for 5 min and the pellet was resuspended with 100 μL sodium dodecyl sulfate (SDS)-loading buffer (100 mM Tris–HCl, pH6.8; 4% SDS; 0.2% bromophenol blue; 20% glycerol; 200 mM β- mercaptoethanol). The whole-cell lysates were subjected to SDS-polyacrylamide gel electrophoresis (SDS-PAGE) and western blot as described using polyclonal rabbit sera raised against the indicated antigens [4].

### Immunization of chickens

Groups of one-day-old Cornish × Rock broiler chickens were purchased from the Murray McMurray Hatchery (Webster City, IA). Chickens were divided into separate pens, with 16-20 chicks per group. On the day of arrival, three chicks were euthanized. Spleen and ceca were collected to confirm their *Salmonella*- free status. The shipping boxes were also swabbed. Tissues were homogenized in PBS and plated onto Brilliant Green plates for bacterial enumeration. Tissue samples and swabs were also enriched in Rappaport-Vassiliadis medium for 48 h at 37°C and subsequently plated onto Brilliant Green. No *Salmonella* were detected in the birds nor in the shipping boxes. The following day (day 0, the birds were 4 days of age), all chicks were orally inoculated with approximately 1 × 10^8^ CFU (experiment 1) or 1×10^9^ CFU (experiments 2 and 3) of various *S*. Typhimurium vaccine strains, including the vector-only control strain χ11802(pYA3681), in 100 μL of PBS. Additional groups of control birds were not vaccinated. Thirty minutes later, the chicks were provided with feed and water. The same dose of the same strain was given as a boost immunization 14 days later.

We performed three independent experiments. Intestinal mucosa samples and cellular response data were not collected in experiment 1 and the strain carrying pKR023 was not tested. In experiment 2, all vaccine strains were used and immune response data were collected, but no challenge was performed. In experiment 3, all strains were used and serum, mucosal and cellular response data were collected.

### Sample collection for analysis of immune responses

To evaluate mucosal immune responses (experiments 2 and 3), intestinal samples were collected as described previously [30] with some modifications. Three birds from each group were necropsied at 19 (experiment 2) or 21 (experiment 3) days after the 1^st^ immunization (6 days after the boost). The intestines were opened aseptically and the surface contents were removed gently using a clean paper wiper. Then, approximately 3 g of intestinal scrapings were collected using glass slides and resuspended in 30 mL of ice-cold PBS containing Pierce™ Protease Inhibitor Mini Tablets. After shaking for 1 min, the supernatants were collected by centrifugation at 4000 × *g* for 20 min at 4°C to evaluate intestinal IgA, IgY and IgM antibody production. Blood was collected from wing veins of all remaining birds in each group at 21 days after the 1^st^ immunization to assess the IgY antibody responses to *C. perfringens* and *Salmonella* antigens in serum.

### Determination of antibody response by enzyme-linked immunosorbent assay (ELISA)

ELISAs were performed in triplicate as described [31] to determine the IgY responses against his-tagged PlcC, his-tagged Fba, his-tagged NetB and *S.* Typhimurium lipopolysaccharide (LPS) in chicken sera and IgA, IgY and IgM responses in intestinal washes. Biotinylated anti-chicken IgA (Alpha Diagnostic Intl. Inc), IgY (Southern Biotechnology) or IgM (Bioss) antibodies diluted 1:10,000 were used to detect the various antibody isotypes.

### Cellular proliferation assay

A proliferation assay was performed to evaluate cell-mediated immunity. Twenty-one days post primary immunization, blood and spleens were harvested. Lymphocytes in the blood were harvested using the gentle swirl technique [32] and plated in quadruplicate, in a 96-well plate at 10^5^ cells/well in RPMI-1640 without phenol red. Spleens were placed through a 70 μm cell strainer to obtain single cell suspensions. Red blood cells were lysed with Red Blood Cell Lysis solution (eBioscience). Splenocytes were then washed, suspended in RPMI and plated at 10^6^ cells/well. Each set of cells was incubated at 37°C, 5% CO_2_ for 72 h with or without 4 μg/ml of either His-Fba, *S*. Typhimurium LPS, His-NetB, His-PlcC, or 1 μg/ml PMA. Cell proliferation was measured using the Vybrant®MTT Cell Proliferation Assay Kit (Molecular Probes). Mean absorbance value of antigen stimulated wells divided by mean absorbance of non-stimulated control wells was used to calculate stimulation index.

### Challenge with *C. perfringens*

Chickens were fed an antibiotic-free starter feed containing 21% protein for 20 days, at which time the feed was switched to a high protein (28% protein), wheat-based feed containing 36% fish meal and zinc at 400 ppm (customized by Reedy Fork Farm, NC) to predispose the birds to necrotic enteritis [11, 33]. Birds were challenged with virulent *C. perfringens* strain CP4 [25] in feed from day 28 to day 32 as described [11] with some modifications. Feed was withdrawn on day 27 for 15 h before challenge. On day 28, chickens were orally gavaged with 0.5 mL of an overnight culture of *C. perfringens* CP4 grown in CMM medium. Immediately after gavage, infected feed was provided thereafter for 5 consecutive days. To prepare infected feed, *C. perfringens* was grown in CMM medium for 24 h at 37°C, which then was inoculated into FTG medium at a ratio of 0.3% (v/v) and incubated at 37°C for 15 h (morning challenge) or 23 h (evening challenge). The *C. perfringens* culture was mixed with feed at a ratio of 1:1 (v/w). Infected feed was prepared freshly twice daily. All birds were euthanized and necropsied the day following the final challenge (day 33).

### Lesion scoring

Protection against *C. perfringens* challenge was assessed on the basis of gross intestinal lesion scores at necropsy. On day 33, chickens were euthanized with CO_2_ and their small intestines (defined here as the section between the gizzard and Meckel’s diverticulum) were examined for visible gross lesions. Intestinal lesions were scored as follows: 0 = no gross lesions; 1 = thin or friable wall or diffuse superficial but removable fibrin; 2 = focal necrosis or ulceration, or non-removable fibrin deposit, 1 to 5 foci; 3 = focal necrosis or ulceration, or non-removable fibrin deposit, 6 to 15 foci; 4 = focal necrosis or ulceration, or non-removable fibrin deposit, 16 or more foci; 5 = patches of necrosis 2 to 3 cm long; 6 = diffuse necrosis typical of field cases [11].

### Attachment assay

The human colon carcinoma cell line Caco-2 (ATCC®#HTB-37) were obtained from the American Type Culture Collection (Manassas, VA) and cultured in Dulbecco’s’ modified Eagle’s medium (DMEM) with 4.5 g/L glucose (Corning, Manassas, VA) containing 4mM L-glutamine, 1% sodium pyruvate, 1% non-essential amino acids (NEEA), 100 U/ml penicillin, 100 μg/ml streptomycin, 20% heat inactivated fetal calf serum. Caco-2 cells were seeded at 5 × 10^5^ cells/mL in each well of a 24-well tissue culture plate. Cells were allowed to grow to a confluent monolayer for 48 hours. One hour prior to infection with CP4, media was replaced with antibiotic free DMEM. Then, 1 μg/ml of purified protein (his-FBA, his-NetB, or his-PspA) or PBS was added to the Caco-2 monolayer and incubated for 15 min at 37°C, 5% CO_2_. The bacterial inoculum was prepared as follows. CP4 cultures were grown anaerobically for 24 h in BHI broth inoculated from a fresh blood agar plate. Cultures were pelleted and resuspended in PBS to a target concentration of 5 × 105 CFU/ 20 μL. Caco-2 monolayers were infected with an MOI of 1:1 and then centrifuged for 3 min at 240 × *g* and incubated for 1 hour at 37°C, 5% CO_2_. To detach the Caco-2 monolayers, 200 μL/well of 0.25% Trypsin-EDTA was added to each well. Once the monolayer cells detached, 800 μL/well of PBS was added. The trypsinized samples were serially diluted and plated onto blood agar plates for *C. perfringens* enumeration. All counts were normalized to the inoculum concentration and presented as percentage of the no protein control.

### Statistical analysis

All statistics were carried out using GraphPad Prism 6.0 (GraphPad Software, San Diego, CA). Antibody titers and adherence data were analyzed using two-way or one-way ANOVA followed by Tukey’s posttest, respectively. Lesion scores were analyzed using a two-tailed Mann-Whitney test. The values were expressed as means ± SEM, and differences were considered significant at *P* < 0.05.

## Acknowledgements

The authors wish to thank Jacquelyn Kilbourne, Larisa Gilley, Nyja Brown, Donovan Leigh, Penelope Roach and Melody Yeh for their expert technical assistance. We thank Dustin McAndrew, Randall Dalbey and the DACT staff at ASU for expert care and husbandry of our research animals.

## Supporting information captions

**Figure S1.** Growth curves of the χ11802 vaccine strains used in this study. Strains were grown with aeration in LB supplemented with 0.1% arabinose and 0.1% mannose. A. Optical density measurements; B. Colony forming units obtained by plating onto LB + arabinose at the indicated times.

---

**Figure S2.** Immunofluorescence of *C. perfringens* strain CP4. Cells were incubated with pre-immune sera (Pre-Bleed), secondary antibody only (2° α) or the indicated antisera with or without prior incubation with the indicated recombinant proteins as outlined in the Materials and Methods section.

